# Regulation of VEGFR2 and AKT signaling by Musashi-2 in lung cancer

**DOI:** 10.1101/2023.03.29.534783

**Authors:** Igor Bychkov, Iuliia Topchu, Petr Makhov, Alexander Kudinov, Jyoti D. Patel, Yanis Boumber

**Affiliations:** Robert H Lurie Comprehensive Cancer Center, Northwestern University, Chicago, IL 60611; Program in Molecular Therapeutics, Fox Chase Cancer Center, Philadelphia, PA 19111; Cardiology Department, University of Illinois in Chicago; address - 840 S. Wood Street Chicago, IL, 60612; Division of Hematology/Oncology, Section of Thoracic Head and Neck Medical Oncology, Feinberg School of Medicine, Northwestern University, Chicago, IL 60611

**Keywords:** Non-small cell lung cancer, Vascular endothelial growth factor receptor-2, Musashi-2, PTEN

## Abstract

Lung cancer is the most frequently diagnosed cancer type and the leading cause of cancer-related deaths worldwide. Non-small cell lung cancer (NSCLC) represents most of the lung cancer. Vascular endothelial growth factor receptor-2 (VEGFR2) is a member of the VEGF family of receptor tyrosine kinase proteins, expressed on both endothelial and tumor cells which is one of the key proteins contributing to cancer development and involved in drug resistance. We previously showed that Musashi-2 (MSI2) RNA-binding protein is associated with NSCLC progression by regulating several signaling pathways relevant to NSCLC. In this study, we performed Reverse Protein Phase Array (RPPA) analysis of murine lung cancer which nominated VEGFR2 protein as strongly positively regulated by MSI2. Next, we validated VEGFR2 protein regulation by MSI2 in several human NSCLC cell line models. Additionally, we found that MSI2 affected AKT signaling via negative *PTEN* mRNA translation regulation. In silico prediction analysis suggested that both VEGFR2 and PTEN mRNAs have predicted binding sites for MSI2. We next performed RNA immunoprecipitation coupled with quantitative PCR which confirmed that MSI2 directly binds to VEGFR2 and PTEN mRNAs, suggesting direct regulation mechanism. Finally, MSI2 expression positively correlated with VEGFR2 and VEGF-A protein levels in human NSCLC samples. We conclude that MSI2/VEGFR2 axis contributes to NSCLC progression and is worth further investigations and therapeutic targeting.

## 1. Introduction

Lung cancer is second in incidence and first in the mortality among all cancer types in recent years based on World Health Organization and American Cancer Society reports [1,2]. Most lung cancer statistics include both small cell lung cancer (SCLC) and non-small cell lung cancer (NSCLC). In general, NSCLC represents about 85% of all lung cancer [3–5]. Lung cancer typically has a_poor prognosis if tumors have disseminated from the primary site[6]. Besides the most common oncogenic driver mutations, *KRAS* and *TP53*, NSCLC is associated with numerous of genetic alterations like *EGFR* mutations*, ALK* translocation, *VEGFR2* and *c-MET* amplification, and others[7,8]. At this time, in addition to chemotherapy, immunotherapy, and targeted therapies, VEGFR2 and VEGF inhibitors show proven efficacy in NSCLC treatment in stage IV disease in combination with chemotherapy and also show promise when combined with immunotherapy [3,9].

Vascular endothelial growth factor (VEGF-A) and its receptors (VEGFR1, VEGFR2, VEGFR3) are the main pro-angiogenic drivers of solid tumors [10]. I the canonical model, VEGF-A is expressed by tumors, which recognizes and binds VEGFR2 on endothelial cells, thus leading to tumor vessel formation[11] and as a result a better supply of tumor cells with nutrients. Additionally, VEGF-A promotes immunosuppressive tumor microenvironment via several mechanisms [12–15]. Published papers show that VEGFR2 and VEGFR1 are well expressed on tumor cells[16–19]. Moreover, *VEGFR2* gain of function is associated with chemoresistance and poor survival of patients with lung cancer[20]. Currently, several drugs are used in the clinic, including VEGFR2-inhibitors (ramucirumab, cabozantinib, pazopanib), VEGF-A inhibitor (bevacizumab)[3], and a VEGF-A/PIGF trap (aflibercept)[21]. Adding VEGFR2 inhibitor, nintedanib, to second-line docetaxel showed improved progression-free survival (median 3.4 months with the combination therapy versus 2.7 months with docetaxel alone; HR 0.79, P=0.0019)[22]. Also, randomized Phase III study compared a combination of ramucirumab with docetaxel to docetaxel alone and reported a modest improvement in median overall survival from 9.1 months to 10.5 months (HR 0.86, P=0.023)[23]

Furthermore, Musashi-2 (MSI2) protein expression is associated with advanced NSCLC [24,25]. MSI2 and its homolog, Musashi-1 (MSI1), are family of RNA-binding proteins that regulate the stability and translation of target mRNAs through high-specific RNA binding of core motif in 3’-untranslated region of mRNAs (UAG) [26–28]. MSI2 regulates multiple critical biological processes in stem cells and cancer cells and contributes to cancer drug resistance [29,30]. MSI2 is upregulated in many hematopoietic and solid tumors, including lung cancer [24,27,31–34]. Our group recently established MSI2 as upregulated in a subset of aggressive NSCLC, and demonstrated a specific role for MSI2 in promoting metastasis in these tumors, through induction of TGFβR1 and its effector SMAD3 and other mechanisms[25]. Furthermore, we have recently shown that MSI2 deficiency leads to higher sensitivity to Epidermal Growth Factor Receptor (EGFR) inhibitors in EGFR mutant NSCLC due to EGFR protein downregulation due to MSI2-mediated direct regulation of EGFR mRNA[24].

In the light of published papers, we have validated proteomic analysis data that suggested VEGFR2 regulation upon MSI2 depletion in murine lung cancer. We found that MSI2 depletion leads to strong decrease in VEGFR2 protein levels in murine and human NSCLC cell lines. Furthermore, we found that MSI2 protein directly binds to *VEGFR2* and *PTEN* mRNAs and impacts VEGFR2 downstream signaling via PTEN regulation. Finally, <u>MSI2</u> protein expression correlated with VEGFR2 and VEGF-A protein levels in NSCLC patient samples.

## 2. Materials and methods

### Cell culture

Human lung cancer cell lines A549 (*KRAS^34G>A^, CDKNA^1_471de1471^*), Calu-1 (*KRAS^34G>T^, TP53^del^*), H23 (*KRAS^34G>T^, TP53^738G>C^*) were obtained from the American Type Culture Collection (ATCC). Human lung cancer cell line Hcc1171 (*KRAS^34G>T^, TP53^740A>T^*), Hcc461 (*KRAS^35G>A^, TP53^445delT^*), were obtained from UT Southwestern Medical Center. Murine NSCLC cell line (344SQ) from Trp53R172HΔG/+KrasLA1/+ mice were previously described [35]. Initial stocks were cryopreserved, and at every 6-month interval, a fresh aliquot of frozen cells was used for the experiments. No additional authentication was performed. All cells were cultured in RPMI 1640 (Gibco, Gaithersburg, MD) supplemented with 10% FBS (Hyclone, Logan, UT), penicillin (100U/ml), streptomycin (100μg/ml), sodium pyruvate (1 mM) and non-essential amino acids (0.1 mM) under conditions indicated in the figure legends. Hypoxic incubations were done with 1% O_2_, 94% N_2_, and 5% CO_2_

### Antibodies and drugs

Anti-MSI2 (#ab76148), anti-VEGF-A (#ab46154) were obtained from Abcam (Cambridge, UK), anti-VEGFR2 (#2479), anti-phVEGFR2 (#3817) anti-β-actin (#3700), anti-phAKT (T308) (#13038), anti-phAKT (S473) (#4060), anti-AKT total (#2920), anti-phGSKαβ (#9331), anti-GSKαβ total (#5676), anti-PTEN (#14642), anti-phERK (#4370), anti-ERK total (4695), anti-ph4EBP(#2855), anti-phP70SK (#9206), anti-P70S6K total (#2708) anti-rabbit HRP-linked (#7074), anti-mouse HRP-linked (#7076) were obtained from Cell Signaling (Danvers, MA). Doxycycline (#HY-N0565), hygromycin (HY-B0490) and puromycin (HY-B1743A) were obtained from MedChemExpress (Monmouth Junction, NJ).

### Vector construction and lentivirus production

To generate stable cell lines with inducible MSI2 knockdowns, self-complementary single-stranded DNA oligos (Supp Table S1) were annealed and cloned into AgeI/EcoR1 sites of Tet-pLKO-puro vector (Addgene plasmid # 21915). MSI2 ORF (NM_138962.2) was amplified by PCR with specific primers and high-fidelity Ex Taq DNA polymerase (Takara Bio USA, Inc., Mountain View, CA) using a cDNA containing human MSI2 obtained from OriGene (Rockville, MD) as a template, and cloned into XbaI/XhoI sites of pLV-CMV-puro vector (a kind gift from Dr. A. Ivanov. All constructs were validated by direct sequencing. To generate stable cell lines with KDR (VEGFR2) overexpression we used pHage-KDR-hygro vector and pHage-hygro vector as negative control. We used plasmids pHage-KDR (#116754) and pHage-puro (#118692) from Addgene with changed resistance gene to hygromycin using a cDNA containing Hygro gene and cloned into ClaI/NdeI and BfuAI/ClaI sites to generate overexpression constructs. pLV-CMV-hygro vector used as a template for Hygro gene. All generated cell lines used in the study are noted in Supp Table S2.

### SiRNA transfections

SiRNAs targeting human MSI2 (Supp Table S3) and nonspecific control pool siRNAs were purchased from Qiagen (Frederick, MD). Cultured cells at 50% confluence were transfected with siRNA at final concentrations of 50 nmol/L using the Lipofectamine RNAiMAX transfection reagent (Thermo Fisher Scientific, Waltham, MA) according to the manufacturer’s instructions.

### Western blot analysis

Cell lysates preparation and Western blot analysis were performed using standard methods as previously described^2^. Bands signals were detected by X-ray films, films were digitized by photo scanner. Image analysis was done using ImageJ (version 1.53e, National Institutes of Health, Bethesda, MD), with signal intensity normalized to β-actin, 3-4 repeats were used for each experiment quantitation analysis. Final data was analyzed in GraphPad Prism by unpaired t-test or ANOVA to determine statistical significance.

### Cell viability assay

To analyze the effects of compounds treatment on proliferation of cells, cells were plated (500 cells/well) in 96-well cell culture plates in complete media. We used increasing concentrations of compounds to calculate IC50 values for each cell line. After 72 hours incubation with compounds, we used supernatant for CellTiter-Blue® assay (Promega, Fitchburg, WI) to measure OD 562 nm. Data was analyzed GraphPad Prism.

### Reverse transcription and qPCR

RNA was extracted using phenol-chloroform based method. RNA concentration and quantity was measured using NanoDrop Lite (cat# ND-LITE ThermoFisher Scientific). First strand cDNA synthesis was performed with iScript cDNA synthesis kit (cat#1708841, Biorad, Califirnia, USA) according to manufacturer’s instructions. The generated cDNA was diluted tenfold and used as a template for qPCR, which was performed with Applied Biosystems QuantStudio 3 system using PowerTrack™ SYBR Green Master Mix (Applied Biosystems). Relative quantification of genes expression was performed using 2^-ΔΔCt^ method, using primers indicated in Supp Table S4.

### ELISA

The concentration of VEGF-A in human lung cancer cell lines was determined by a VEGF Human ELISA kit (Abcam, #ab100662). ELISA assays were performed according to the manufacturer manual. Briefly, the collected condition media from cells was added to a well coated with primary antibody, and then immunosorbented by biotinylated primary antibody at room temperature for 2.5 h. The color development catalyzed by horseradish peroxidase was terminated with 2.5 mol/l sulfuric acid and the absorption was measured at 450 nm. The protein concentration was determined by comparing the relative absorbance of the samples with the standards.

### RNA-IP assays

RNA was immunoprecipitated from cell lysates (2 × 10^7^ cells per IP) using either a control normal rabbit IgG or rabbit monoclonal anti-MSI2 antibody and the Magna RIP RNA-binding Protein Immunoprecipitation kit (cat#17-700, Millipore, Burlington, MA). Manufacturer’s instructions were followed with the exception that RNeasy MinElute Cleanup kit (cat#74202, Qiagen, Venlo, Netherlands) was used to prepare RNA. Immunoprecipitated RNAs were quantified by quantitative PCR (qPCR) using primers indicated in Supp Table S4, using PTP4A1 as a normalization (positive) control and GAPDH as a negative control.

### RPPA

The 344SQ-SCR, 344SQ-m1, and 344SQ-m2 mouse cells were previously described [25]. Prior to analysis, these cells were lysed and prepared according to MD Anderson Core Facility instructions[26,27,36–39], and RPPA was performed at MD Anderson facility and previously published [25]. Data were visualized using the MultiExperiment Viewer program (www.tm4.org/mev.html) [39].

### Immunohistochemistry of human NSCLC

Surgical NSCLC specimens of patients from the Rostov Research Institute Human Tissue Repository Facility (HTRF) and from Republican Clinical Oncology Dispensary named after prof. M. Z. Sigal (RCOD at Kazan) were used. At the time of tissue acquisition, patients provided Institutional Review Board (IRB)–approved informed consent for storing tissue and reviewing deidentified clinical data. Clinical information (Supp Table S5) from the repository database was abstracted in an anonymized fashion. Tissue samples were stained for VEGFR2 and MSI2 proteins via immunohistochemical (IHC) approach and hematoxylin and eosin (H&E) stained sections were used for morphological evaluation purposes, and unstained sections were used for IHC staining using standard methods. Briefly, 5 μm formalin-fixed, paraffin-embedded sections were deparaffinized and hydrated. Sections were then subjected to heat-induced epitope retrieval with 0.01M citrate buffer (pH 6.0) (MSI2) or EDTA buffer (VEGFR2). Endogenous peroxidases were quenched by the immersion of the slides in 3% H_2_O_2_ solution. The sections were incubated overnight with primary antibodies to MSI2 (EP1305Y, Rabbit, 1:100, Abcam #ab76148), VEGF-A (VG-1, Mouse, 1:50, Abcam #ab1316), VEGFR2 (55B11, Rabbit, 1:50, Cell signaling, Cat #2479) at 4 °C in a humidified slide chamber. As a negative control, the primary antibody was replaced with normal mouse/rabbit IgG to confirm absence of specific staining. Immunodetection was performed using the Dako Envision+ polymer system and immunostaining was visualized with the chromogen 3, 3’-diaminobenzidine. All slides were viewed with a Nikon Eclipse 50i microscope and photo-micrographs were taken with an attached Nikon DS-Fi1 camera (Melville, NY, USA).

### In silico evaluation of MSI2 binding to VEGFR2, VEGF-A and PTEN mRNAs

Human and murine genome sequences for EGFR were obtained from the UCSC Human Gene Sorter December 2013 (GRCh38/hg38) assembly and scanned for Musashi binding motifs previously defined by Bennett et al. [26] (15 motifs with highest p values) and Wang et al. [27] (8 motifs with highest p values; Supp Tables S6 and S7).

### Statistical analysis

All statistical analyses, including unpaired two-tailed t-test, ANOVA analysis, Spearman correlation, were performed in GraphPad Prizm 9 (San Diego, CA).

## 3. Results

### 3.1. Musashi-2 regulates VEGFR2 mRNA and protein levels and directly binds *VEGFR2* mRNA in human NSCLC cell lines

Our published paper suggested that MSI2 depletion in murine lung cancer cell line 344SQ may decrease VEGFR2 protein level based on data (Supplementary Fig. 1A). To verify this result, we used our previously established KRAS/P53 driven 344SQ cell line with doxycycline (DOX) – inducible knockdown (KD) of MSI2 (sh1, sh2) and with constant MSI2 overexpression (OE) (MSI2) (Supplementary Fig. 1B) and evaluated VEGFR2 and VEGF-A protein and mRNA levels (Supplementary Fig. 1C and D). We found that MSI2 KD leads to significant decrease of VEGFR2 protein level, but not mRNA in mouse cells. In addition, MSI2 OE leads to mild increase in VEGF-A mRNA and protein levels, but not VEGFR2 protein and mRNA levels.

Thus, we conclude that MSI2 may regulate VEGFR2 and VEGF-A in mouse cells. Next, based on published papers[19,40,41] we selected several NSCLC cell lines with significant VEGFR2 expression and examined VEGFR2 and VEGF-A protein levels by western blot analysis (Fig. 1A) and *VEGFR2* mRNA level (Fig. 1B). We selected cell lines with higher expression of VEGFR2 (H441, Hcc1171, and Calu-1) and A549 cell line as a negative control / low VEGFR2 expression for further analysis. Previously, Bennett et al.[26], Wang et. al[27], and Nguyen et al.[28] in their papers showed that MSI2 recognizes and directly binds consensus sequences with core motif UAG in 3’-untranslated region (3’-UTR) of mRNAs, and as a result regulates the stability and/or translation of target mRNAs. Based on this published data we performed *in silico* analysis (using Bennett and Wang data) and found that *VEGFR2* and *VEGF-A* mRNAs have predicted binding sites for MSI2 binding (Fig. 1C, Supplementary Tab. 6, 7). To confirm the *in silico* results, we performed RNA immunoprecipitation assays (RIP) with an MSI2 antibody pulldown, coupled with RT-qPCR in three NSCLC cell lines: A549 (Supplementary Fig. 2A), Hcc1171 H441(Fig. 1D), using previously defined MSI2 target mRNAs (*PTP4A, TGFβR1, SMAD3*)[25,42] as positive controls and *ACTB* and *GAPDH* as negative controls. Antibodies to MSI2 specifically immunoprecipitated the *VEGFR2* mRNA as efficiently as they did the positive controls (Fig. 1D). Taken together, we conclude that MSI2 directly regulates VEGFR2 mRNA translation in human NSCLC.

**Figure 1.**
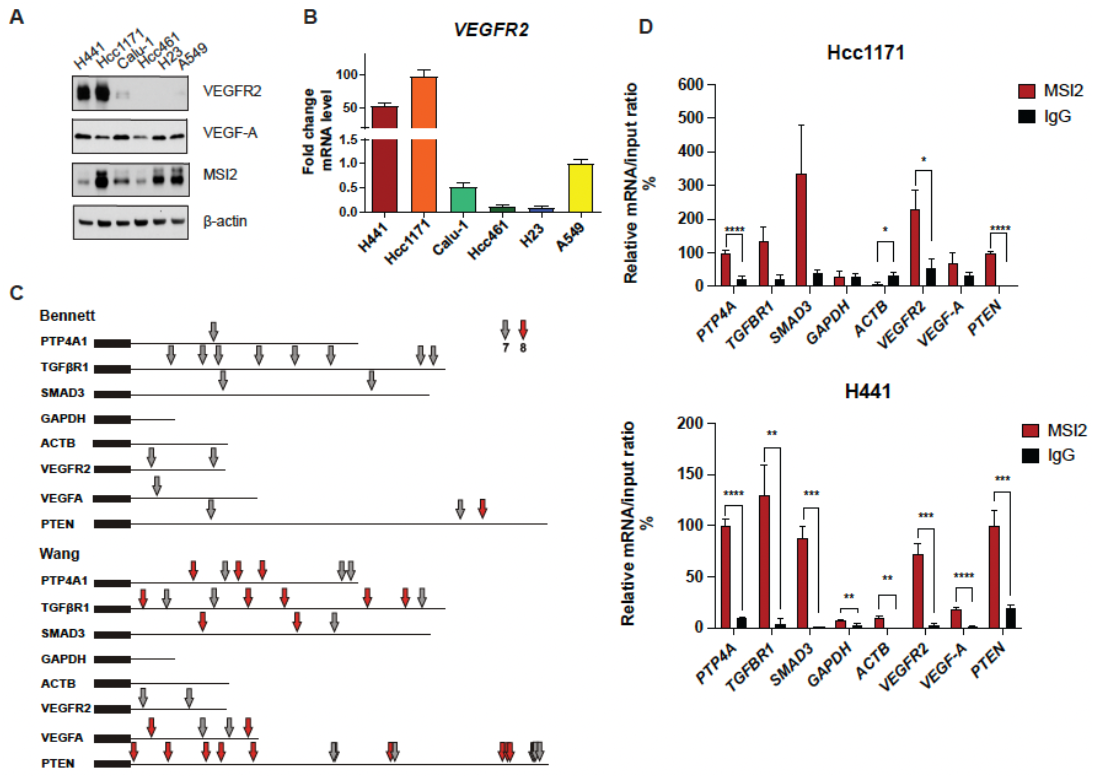
MSI2 directly binds VEGFR2 mRNA in human NSCLC. **(A)** Western blot of human NSCLC cell lines. **(B)** RT-qPCR analysis of VEGFR2 mRNA from human NSCLC cell lines. Data from at least three independent experiments by Image J software and normalized to 18S rRNA and to A549. **(C)** MSI2 consensus binding sites in human mRNAs. Location of consensus binding sites for Musashi proteins in the noted human genes, as defined from studies by Bennett et al^1^, and Wang et al^2^. Coding sequences are represented by thick lines; 3’ untranslated regions by a thin line. 7- or 8-bp consensus sequences are indicated by arrows. *VEGFR2* reference sequence - NCBI Reference Sequence: NM_002253.4, *PTEN* reference sequence - NCBI Reference Sequence: NM_000314.8. **(D)** Quantification of mRNA immunoprecipitation (RIP) results from assays performed in Hcc1171 and H441 cell lysates using antibodies to MSI2, or IgG (negative control) antibodies, followed by quantitative RT-PCR. Data are normalized to positive control *PTP4A1, TGFBR1*, and *SMAD3* are additional positive controls; *GAPDH* and *ACTB* are negative control. The data shown reflect the average of three independent RIP experiments. Error bars represented by SEM. Statistical analysis was performed using an unpaired two-tailed t-test. *p < 0.05, **p < 0.01, ***p < 0.001 for all graphs.

### 3.2. Musashi-2 regulation of VEGFR2 and VEGF-A protein levels in NSCLC

Since MSI2 directly binds *VEGFR2* mRNA in human NSCLC cell lines, we established cell lines with MSI2 depletion and overexpression to evaluate <u>the</u> effect of MSI2 expression on VEGFR2 and VEGF-A protein levels. Depletion of MSI2 in human NSCLC cell lines has led to significant decrease of VEGFR2 protein level and mixed effect of intracellular VEGF-A protein level (Fig. 2A and B, Supplementary Fig. 2B). Also, MSI2 OE leads to slight increase of VEGFR2 protein level in A549 and H441 cell lines and significant decrease of VEFR2 protein level in Hcc1171 and Calu-1 cell lines (Supplementary Fig. 2 C and D). In addition, we performed ELISA analysis of extracellular VEGF-A concentration after MSI2 depletion and overexpression (Fig. 2 C). ELISA analysis indicated that MSI2 depletion leads to VEGF-A decrease only in A549 cell line, while MSI2 OE leads to significant increase of VEGF-A concentration in cell lines that have moderate to high expression of VEGFR2 (H441, Hcc1171 and Calu-1). Moreover, RT-qPCR analysis of *VEGFR2* and *VEGF-A* with MSI2 depletion and overexpression showed that *VEGFR2* mRNA is significantly decreased in human NSCLC cell lines with MSI2 depletion and increased with MSI2 OE in cell lines with moderate to high VEGFR2 expression (H441, Hcc1171, and Calu-1) (Supplementary Fig.3). In parallel, evaluation of MSI2 depletion effect on cell growth in these cell lines showed that MSI2 level doesn’t affect NSCLC cell growth (Supplementary Fig. 4A). Therefore, we conclude that MSI2 positively regulates both *VEGFR2* mRNA and protein levels and it also regulates *VEGF-A* mRNA translation in human lung cancer cells.

**Figure 2.**
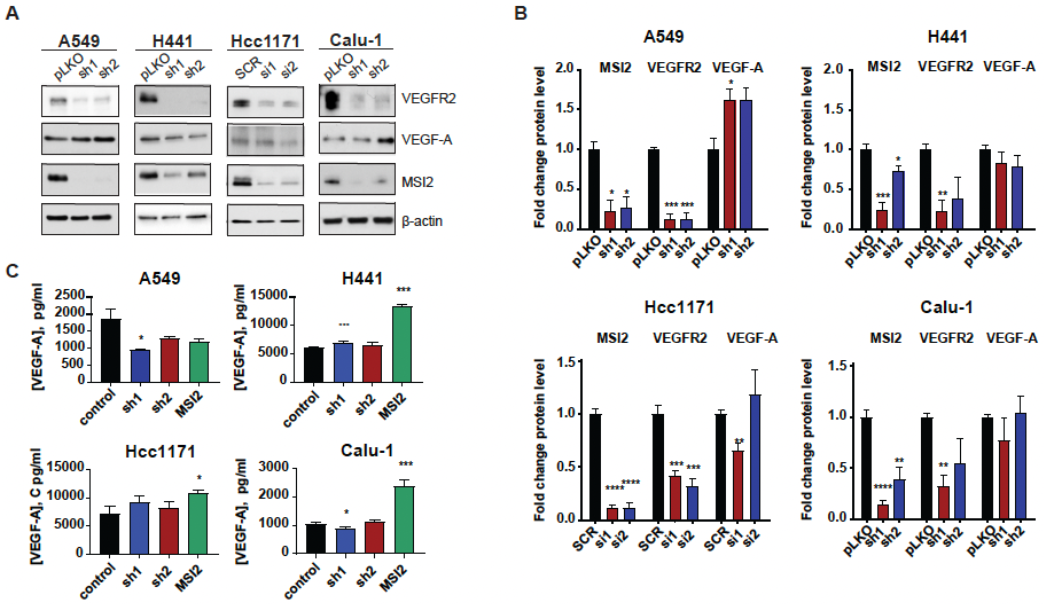
MSI2 regulation of VEGFR2 and VEGF-A protein levels in human NSCLC cell lines. **(A)** Western blots of indicated cell lines, following depletion by shRNA (sh1, sh2) and siRNA (si1, si2) of MSI2. Negative controls include pLKO and SCR. MSI2 depletion was induced by the addition of 1 μg/ml of Doxycycline for 48 h. **(B)** Quantification of Western blot (Fig 2A) data from at least three independent experiments by Image J software, with values normalized to negative control and β-actin. **(C)** The concentration of VEGF-A in cell culture media from indicated cell lines, following depletion by shRNA (sh1, sh2), siRNA (si1, si2) and overexpression (MSI2) of MSI2. MSI2 depletion was induced by the addition of 1 μg/ml of Doxycycline for 48 h. The ELISA data shown reflects the average of three independent experiments. Error bars represented by SEM. Statistical analysis was performed using an unpaired two-tailed t-test. *p < 0.05, **p < 0.01, ***p < 0.001 for all graphs.

### 3.3. Musashi-2 regulates AKT signaling via *PTEN* mRNA binding independent of VEGFR2

Since MSI2 depletion leads to reduction of VEGFR2 protein levels, we evaluated VEGFR2 downstream signaling in human NSCLC cell lines. Western blot analysis showed that MSI2 depletion reduces VEGFR2 and phAKT protein levels (Fig. 3 A and B, Supplementary Fig. 4B). To assess the role of VEGFR2 in downregulation of phAKT with MSI2 KD, we established A549 and H441 cell lines with inducible MSI2 KD and constant VEGFR2 overexpression (OE). Analysis of AKT signaling indicated that MSI2 KD leads to decrease of phAKT and its downstream target phGSK3α/β with and without VEGFR2 OE of VEGFR2 (Fig. 3 C and D). These findings suggest that decrease in VEGFR2 protein level doesn’t affect AKT signaling.

**Figure 3.**
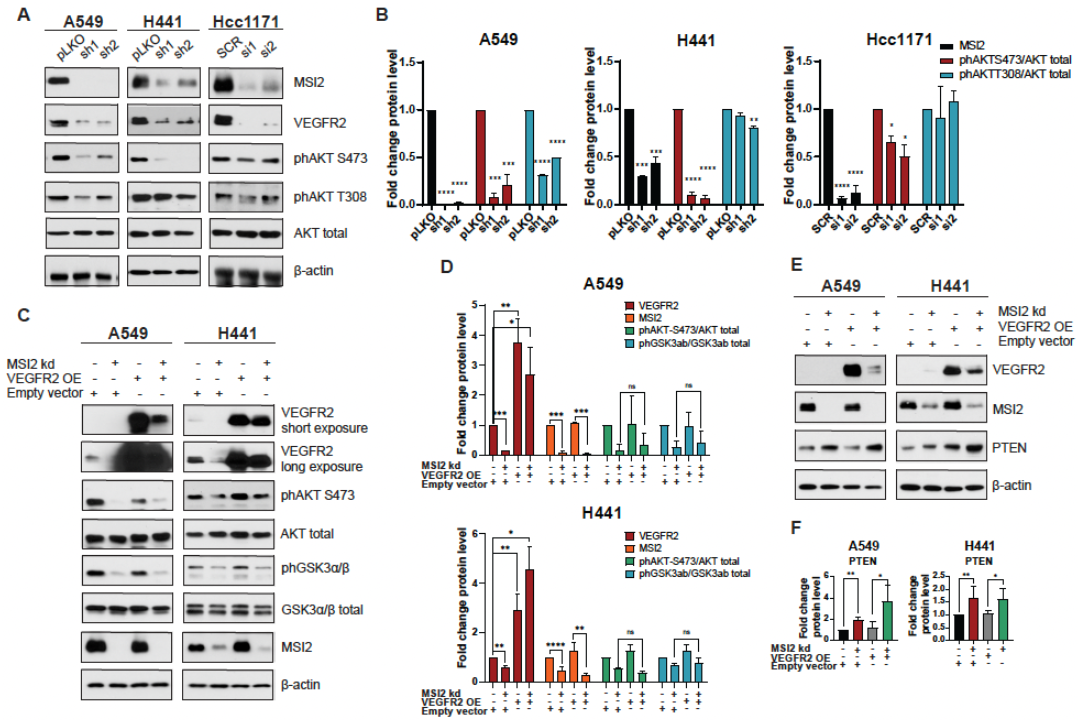
VEGFR2-independent effects of MSI2 KD on AKT signaling. **(A)** Western blots of indicated cell lines, following depletion by shRNA (sh1, sh2) and siRNA (si1, si2) of MSI2. Negative controls include pLKO and SCR. **(B)** Quantification of Western blot data (Fig. 3A) from at least three independent experiments by Image J software, with values normalized to negative control and β-actin. **(C)** Rescue experiment: western blots of A549 and H441 cell lines, following depletion by shRNA (sh1) of MSI2 and VEGFR2 overexpression (VEGFR2 OE). Negative control is pHAGE. **(D)** Quantification of Western blot data (Fig. 3C) from at least three independent experiments by Image J software, with values normalized to empty vector and β-actin. **(E)** Western blot of indicated cell lines, following depletion by shRNA (sh1) of MSI2 and VEGFR2 overexpression (VEGFR2 OE). **(F)** Quantification of Western blot data (Fig. 3E) from at least three independent experiments by Image J software, with values normalized to empty vector and β-actin. Error bars represented by SEM. Statistical analysis was performed using an unpaired two-tailed t-test. *p < 0.05, **p < 0.01, ***p < 0.001, ****p <0.0001.

Wang et al.[27] showed that MSI2 inhibits PTEN (tumor suppressor and inhibitor of AKT signaling) protein level in murine intestinal epithelium. Therefore, we evaluated PTEN protein level in human NSCLC cell lines upon MSI2 depletion (Fig. 3E and F). Western blot analysis showed significant upregulation of PTEN with MSI2 KD regardless of VEGFR2 level. This suggests that MSI2 may directly regulate *PTEN* mRNA level and affect AKT signaling in NSCLC. To test this, we performed *in silico* analysis and found that 3’ UTR of *PTEN* mRNA has predicted binding sites for MSI2 (Fig.1D, Supplementary Fig. 2A). Next, we performed RIP-QPCR analysis using MSI2 antibody which supported the *in-silico* predictions and showed that MSI2 directly binds *PTEN* mRNA. Taken together, we conclude that MSI2 affects AKT signaling via direct *PTEN* mRNA binding, independent of *VEGFR2* mRNA regulation.

### 3.4. Correlation of MSI2 with VEGFR2 and VEGF-A expression in human NSCLC

To evaluate the relationship between MSI2, VEGFR2 and VEGF-A in the subset of NSCLC tumors, we performed IHC analysis of MSI2, VEGFR2 and VEGF-A expression in an independent group of 147 NSCLC clinical tumor samples (Fig. 4A and B). Spearman’s analysis of H-scores showed moderate positive correlation MSI2 vs VEGFR2 (r = 0.544) and MSI2 vs VEGF-A (r = 0.548).

**Figure 4.**
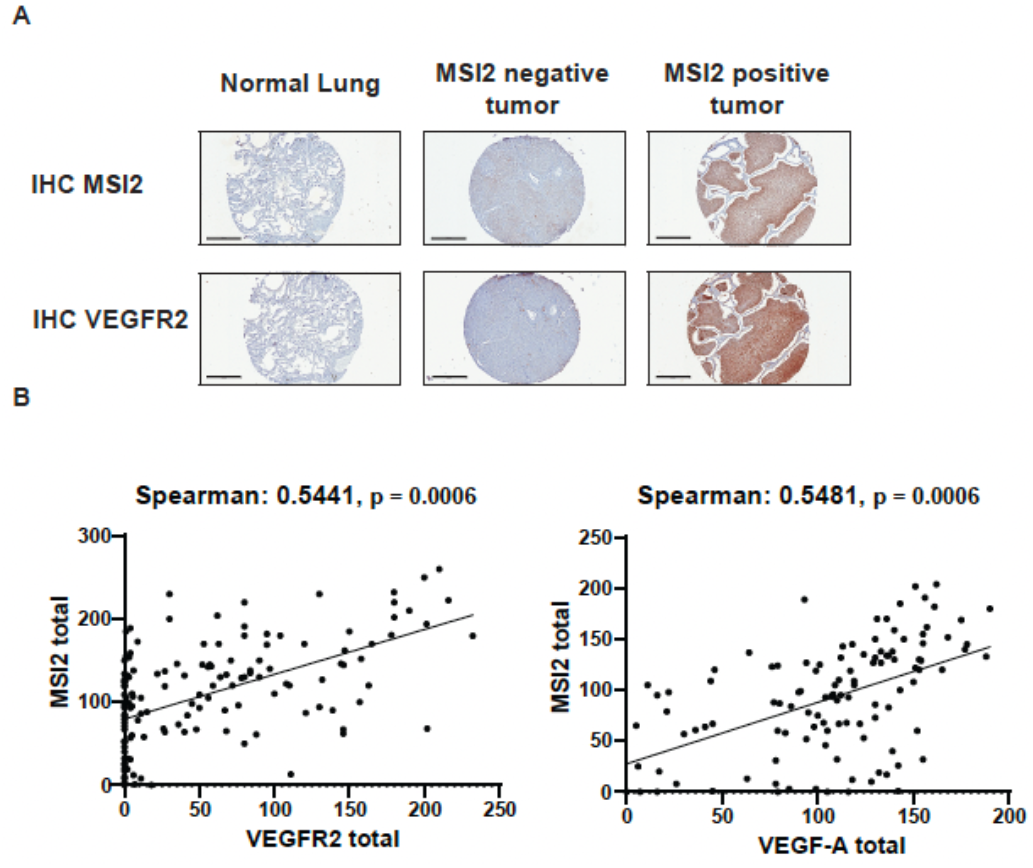
Expression of MSI2, VEGFR2 and VEGF-A proteins in human NSCLC patient samples. **(A)** Representative IHC images of MSI2 and VEGFR2 expression in normal lung and lung tumors. **(B)** MSI2 and VEGF-A (n=94), MSI2 and VEGFR2 (n=118) H-score correlation of human NSCLC TMAs. For IHC quantification, each spot was examined by board-certified pathologists (ED and NK) who assigned a score of 0 (no staining), 1+ (weak staining), 2+ (moderate staining), and 3+ (strong staining) within carcinomatous areas. The score for each of the two tumor spots was averaged for statistical analysis. The H-score, which ranges from 0 to 300, was calculated using the following formula: [1x(% cells 1+) + 2x(% cells 2+) + 3x(% cells 3+)], which reflects staining intensity as well as percentage of positive cells. A sum of p2 and p3 represents a sum of 2+ and 3+ cells (2x (% cells 2+) + 3x(% cells 3+), which excludes 1+ cells.

## 4. Discussion

Published papers show that VEGFR2 and VEGF-A proteins are widely expressed by various tumor types, including lung [17–19,43]. Additionally, *VEGFR2* overexpression is associated with chemoresistance and poor survival of patients with lung cancer [20]. Our study for the first time shows that MSI2 positively regulates VEGFR2 protein level in NSCLC.

First, we selected human lung NSCLC cell lines with high levels of *VEGFR2* expression (H441, Hcc1171 and Calu-1) and A549 as a control with low level VEGFR2 (Fig. 1A and B). MSI2 typically regulates its targets by direct binding to specific motifs in 3’-UTR fragments of mRNAs [26–28]. Therefore, we performed *in silico* analysis and then validated it by RNA-IP analysis (Fig. 1C and D). *In silico* analysis and RNA-IP validation showed that MSI2 directly binds and *VEGFR2* but not *VEGF-A* mRNA. These results expand our knowledge about the role of Musashi-2 in NSCLC progression.

Then, we established human NSCLC cell lines with MSI2 KD and MSI2 OE to see how MSI2 level effects VEGFR2 and VEGF-A protein and mRNA levels (Fig. 2, Supplementary Fig. 3). We found that MSI2 depletion leads not only to decrease of VEGFR2 protein level, but also mRNA level. That result can be interpreted in a way that MSI2 regulates not only protein translation, but also affects mRNA transcript stability[29]. Moreover, we found that MSI2 overexpression leads to modest increase of extracellular VEGF-A from cells with *VEGFR2* overexpression. This effect may be interpreted as indirect positive regulation of VEGF-A by MSI2[44,45].

As MSI2 regulates VEGFR2 protein level, we evaluated cell proliferation along with VEGFR2 downstream signaling in human NSCLC cell lines upon MSI2 depletion. Viability assay showed that MSI2 doesn’t affect cell growth in these KRAS-driven cells (Supplementary Fig. 4A). Signaling evaluation indicated decrease VEGFR2 and phAKT protein levels with MSI2 KD (Fig. 3A and B), while phERK protein level not changed (Supplementary Fig.4B). This result is expected, because our models are *KRASmut*, and published papers show that *KRASmut* results in constitutive activation of ERK[46,47]. Next, we established cell lines with VEGFR2 OE to evaluate MSI2 effects AKT signaling via VEGFR2 regulation. Our results indicated that MSI2 KD induces decrease of phAKT and its downstream target GSK3α/β regardless of the VEGFR2 level (Fig. 3C and D).

Wang et al.[27] previously showed that MSI2 inhibits PTEN protein level in murine intestinal epithelium. Based on that, evaluation of PTEN level with MSI2 KD and VEGFR2 OE indicated that PTEN protein levels are increased upon MSI2 KD regardless of the VEGFR2 level (Fig. 3E and F). In addition, we showed that MSI2 negatively directly regulates PTEN protein level (Fig. 1C and D, Supplementary Fig. 2A). We next performed in silico analysis which predicted MSI2 binding sites in PTEN mRNA, and such binding was validated using RNA IP / QPCR analysis. Taken together, our data suggests that PTEN is direct MSI2 target in NSCLC. Therefore, MSI2 affects AKT signaling via direct *PTEN* mRNA binding, independent of *VEGFR2* mRNA regulation.

In addition, TMA analysis of human lung tumor samples showed positive correlation between MSI2 and VEGFR2 and VEGF-A protein levels (Fig. 4). Taken together, we conclude that previously unsuspected novel Musashi-2 / VEGFR2 signaling axis is worth additional investigations and could be targeted for better NSCLC control in the future.

## Acknowledgments

We acknowledge assistance from the Fox Chase Cancer Center Cell Culture Facility, Histopathology Facility, Biostatistics Facility, and Biosample Repository. This work and the authors were supported by NIH R01 CA218802 grant (to Y.B.); Translational Bridge Award from Northwestern University number 2022-001 (to Y.B.); the NCI Core Grant P30 CA060553 (to Robert H Lurie Comprehensive Cancer Center at Northwestern University); NCI R21 CA263362 grant (to P.M.).

---

**Supplementary Figure 1.**
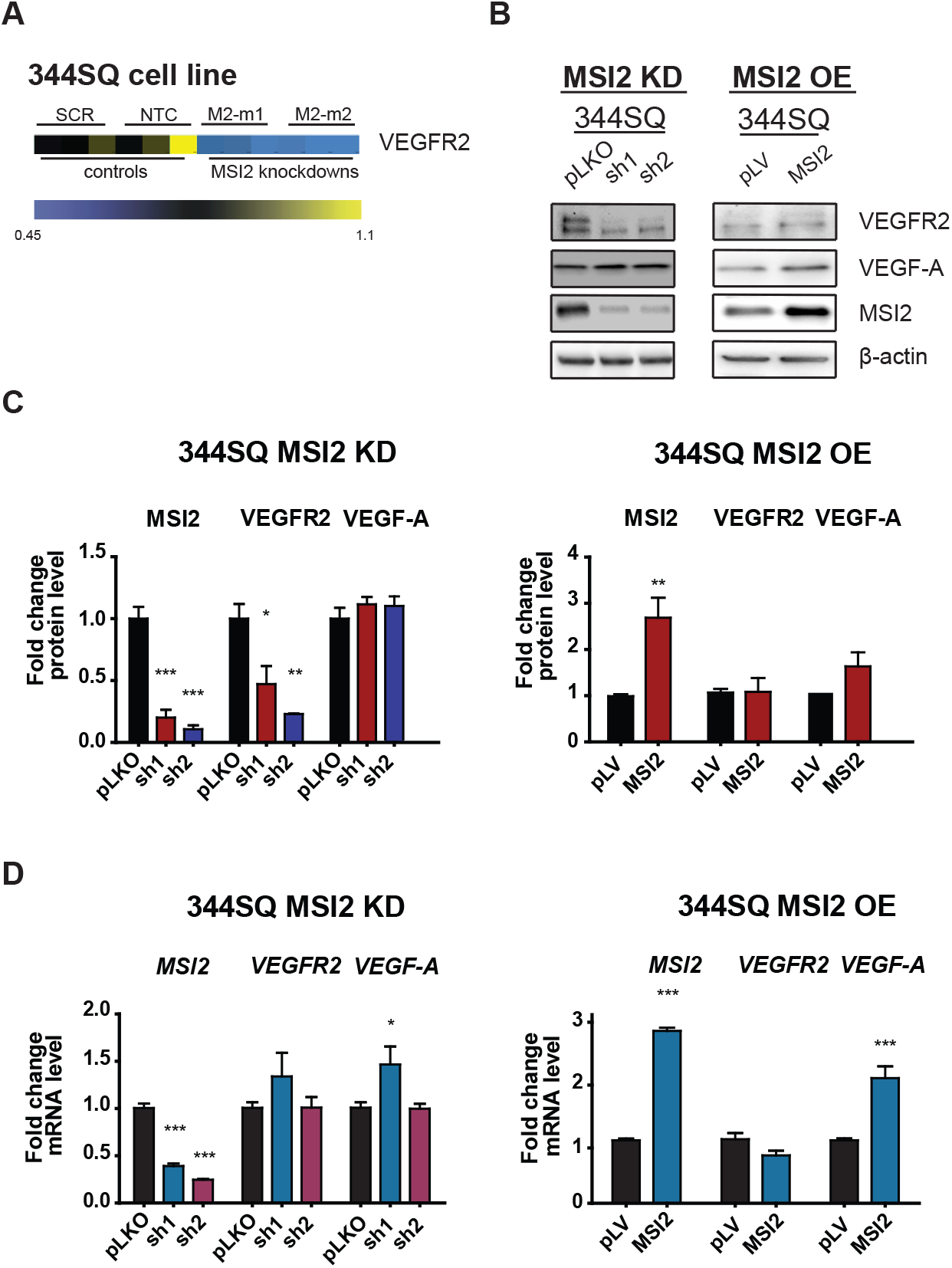
MSI2 regulation of VEGFR2 protein level in a mouse cell line. (A) Heatmap of RPPA results for VEGFR2 protein expression in 344SQ cell line transfected with empty vector or shRNAs or MSI2 depletion with two independent shRNAs (M2-m1 and M2-m2). (B) Western blots of 344SQ cell line lysates, following depletion (MSI2 KD sh1, sh2) and overexpression (MSI2 OE, MSI2) of MSI2. pLKO and pLV are negative controls. MSI2 depletion was induced by the addition of 1 μg/ml of Doxycycline for 48 h. (C) Quantification of Western blot data (Supplementary Fig. 1A) from at least three independent experiments by Image J software, with values normalized to negative control and β-actin. (D) RT-qPCR analysis of Vegfr2 and Vegf-a mRNA from 344SQ cell line following depletion (MSI2 KD, sh1, sh2) and overexpression (MSI2 OE, MSI2) of MSI2. pLKO and pLV are negative controls. MSI2 depletion was induced by the addition of 1 μg/ml of Doxycycline for 48 h. Data normalized to negative control and Polr2a. Error bars represented by SEM. Statistical analysis was performed using an unpaired two-tailed t-test. *p < 0.05, **p < 0.01, ***p < 0.001 for all graphs.

**Supplementary Figure 2.**
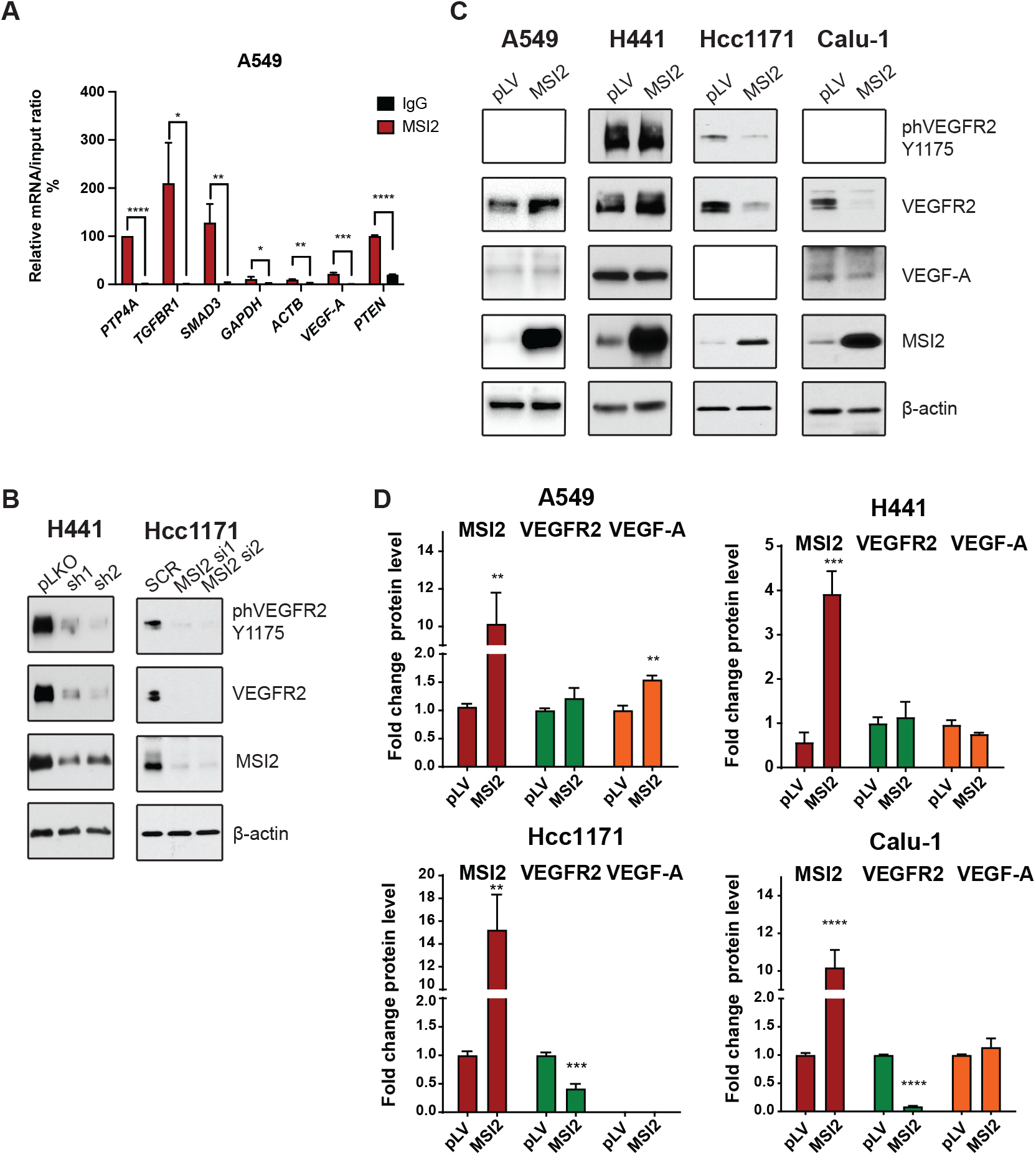
(A) Quantification of mRNA immunoprecipitation (RIP) results from assays performed in A549 cell lysates using antibodies to MSI2, or IgG (negative control) antibodies, followed by quantitative RT-PCR. Data are normalized to positive control PTP4A1, TGFBR1, and SMAD3 are additional positive controls; GAPDH and ACTB are a negative control. The data shown reflect the average of three independent RIP experiments. (B) Western blots of indicated cell lines, following depletion by shRNA (sh1, sh2) and siRNA (si1, si2) of MSI2. Negative controls include pLKO and SCR. (C) Western blot of human NSCLC cell lines, following MSI2 overexpression (MSI2). Negative control is empty vector pLV. (D) Quantification of Western blot data (Supplementary Fig. 2C) from at least three independent experiments by Image J software, with values normalized to negative control and β-actin. Error bars represented by SEM. Statistical analysis was performed using an unpaired two-tailed t-test. *p < 0.05, **p < 0.01, ***p < 0.001.

**Supplementary Figure 3.**
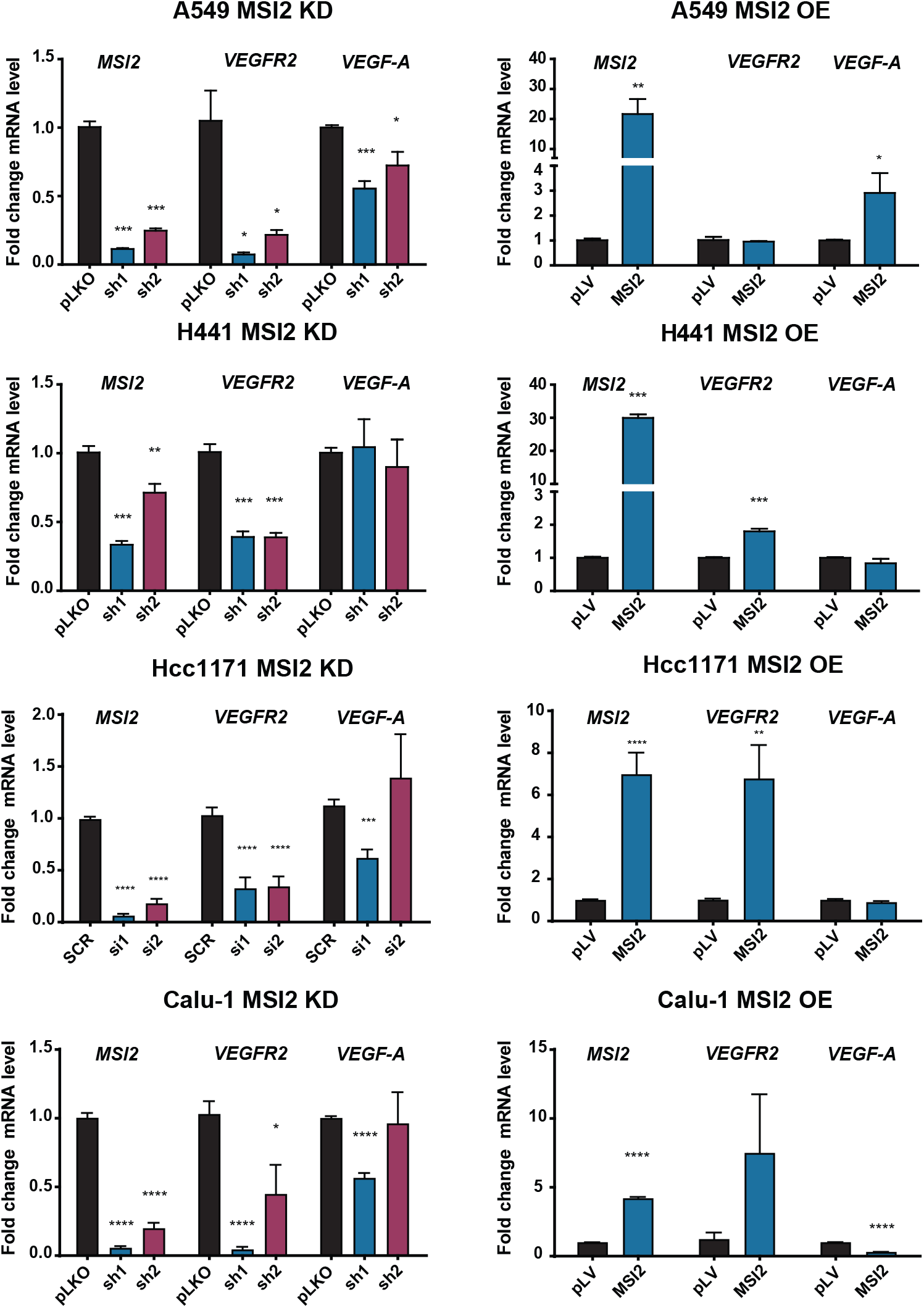
Consequences of MSI2 depletion (left) and overexpression (right) on mRNA expression levels of VEGFR2 and VEGF-A. Quantitative RT-PCR of mRNA of human NSCLC cell lines, following MSI2 depletion by shRNA (sh1, sh2) and siRNA (si1, si2) and overexpression (MSI2) of MSI2. Negative controls include pLKO, SCR, pLV. Error bars represented by SEM. Statistical analysis was performed using an unpaired two-tailed t-test. *p < 0.05, **p < 0.01, ***p < 0.001.

**Supplementary Figure 4.**
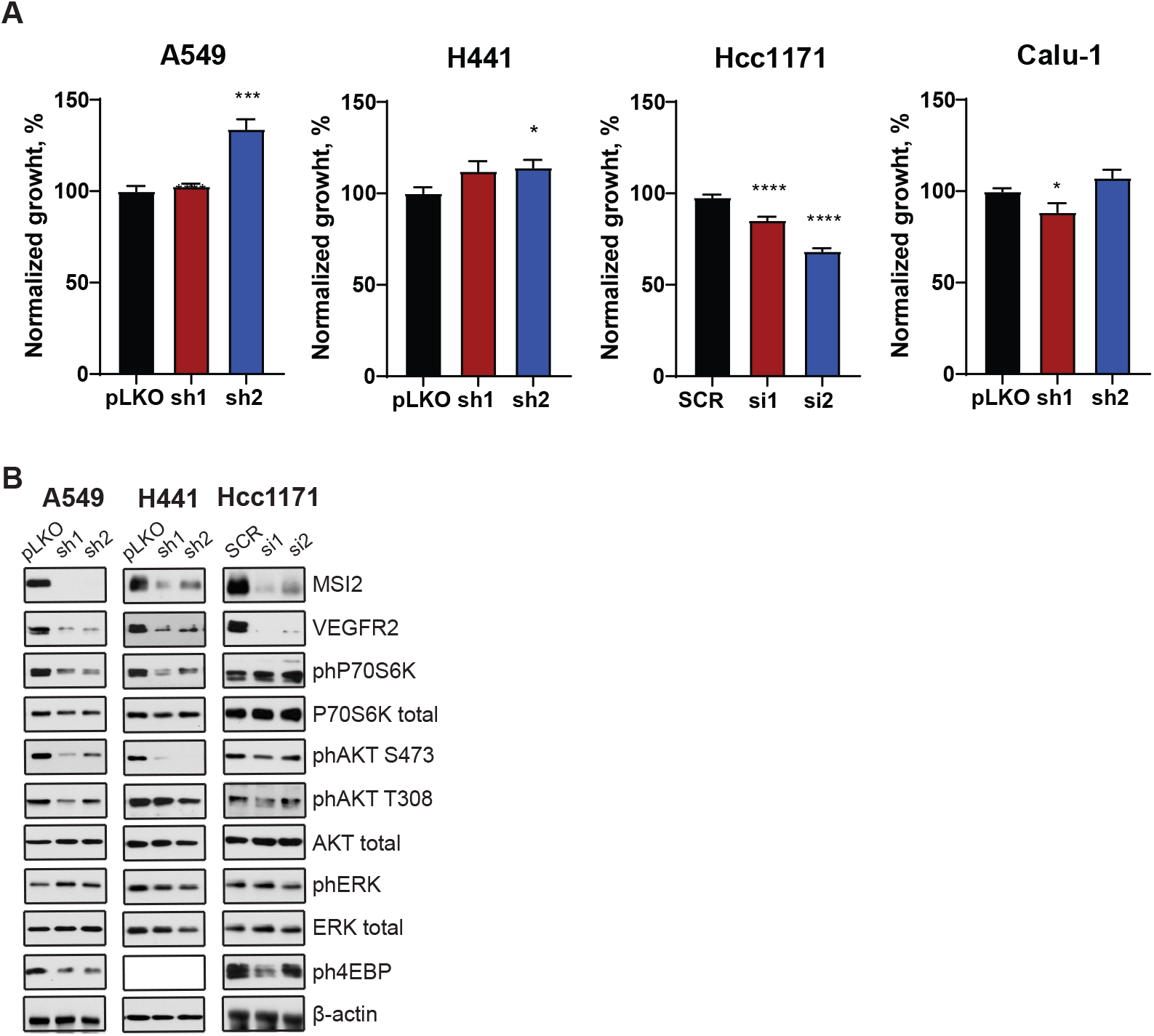
(A) Cell viability quantified by Cell Titer Blue (CTB) assay of indicated cell lines following depletion by shRNA (sh1, sh2), siRNA (si1, si2) and overexpression (MSI2) of MSI2. MSI2 depletion was induced by the addition of 1 μg/ml of Doxycycline for 48h. Negative controls include pLKO or SCR. Error bars represented by SEM. Statistical analysis was performed using an unpaired two-tailed t-test. *p < 0.05, **p < 0.01, ***p < 0.001, ****p <0.0001. (B) Western blots of indicated cell lines, following depletion by shRNA (sh1, sh2) and siRNA (si1, si2) of MSI2. Negative controls include pLKO and SCR. Error bars represented by SEM. Statistical analysis was performed using an ANOVA. *p < 0.05, **p < 0.01, ***p < 0.001, ****p <0.0001.

