## Supplementary material for "Regulation of VEGFR2 and AKT signaling by Musashi-2 in lung cancer": figS2.pdf

### Supplementary Figure 2

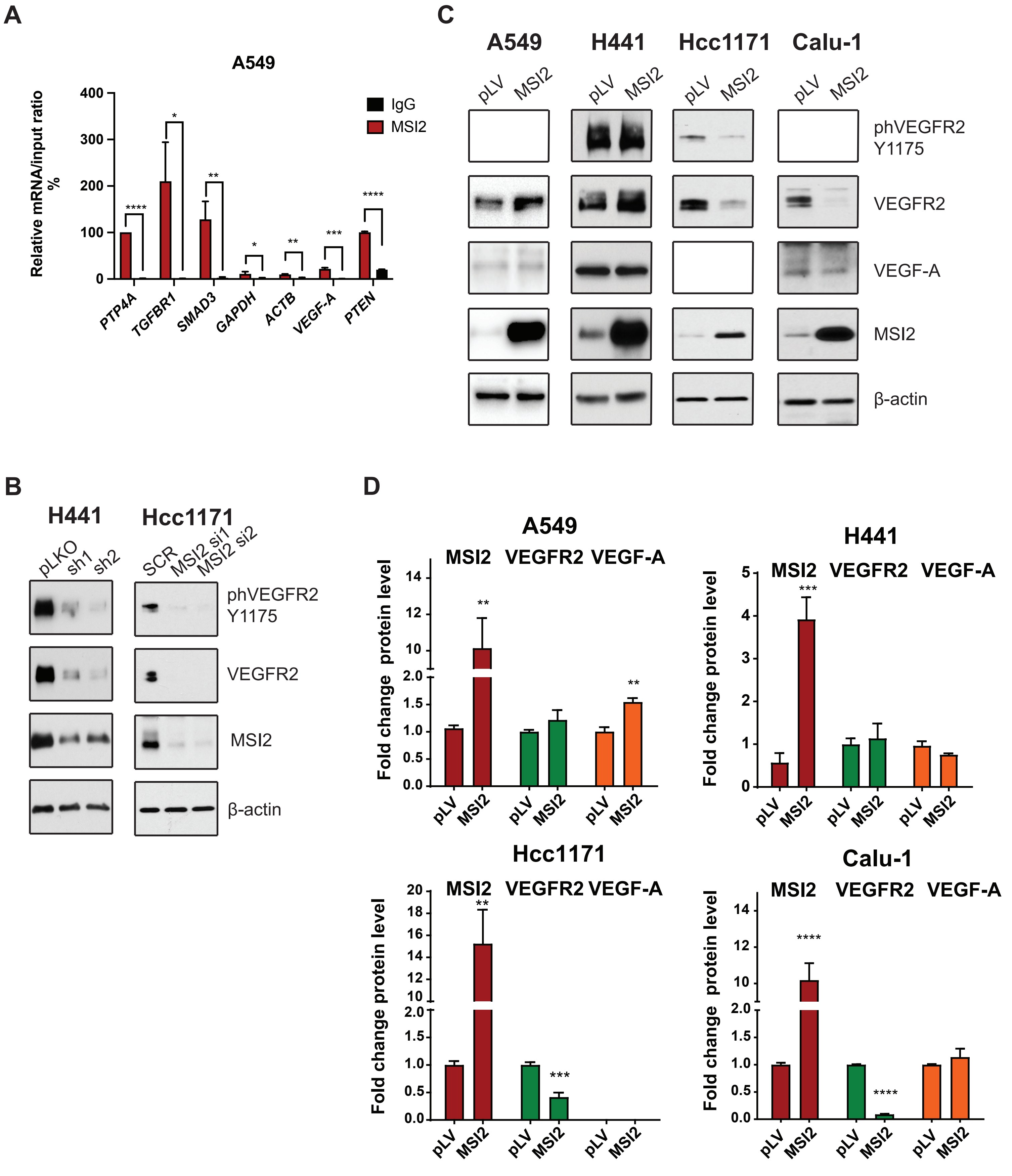


Supplementary Figure 2. (A) Quantification of mRNA immunoprecipitation (RIP) results from assays performed in A549 cell lysates using antibodies to MSI2, or IgG (negative control) antibodies, followed by quantitative RT-PCR. Data are normalized to positive control PTP4A1, TGFBR1, and SMAD3 are additional positive controls; GAPDH and ACTB are a negative control. The data shown reflect the average of three independent RIP experiments. (B) Western blots of indicated cell lines, following depletion by shRNA (sh1, sh2) and siRNA (si1, si2) of MSI2. Negative controls include pLKO and SCR. (C) Western blot of human NSCLC cell lines, following MSI2 overexpression (MSI2). Negative control is empty vector pLV. (D) Quantification of Western blot data (Supplementary Fig. 2C) from at least three independent experiments by Image J software, with values normalized to negative control and β-actin. Error bars represented by SEM. Statistical analysis was performed using an unpaired two-tailed t-test. *$p < 0.05$, **$p < 0.01$, ***$p < 0.001$.
