## Supplementary material for "Regulation of VEGFR2 and AKT signaling by Musashi-2 in lung cancer": figureS3.pdf

### Supplementary Figure 3

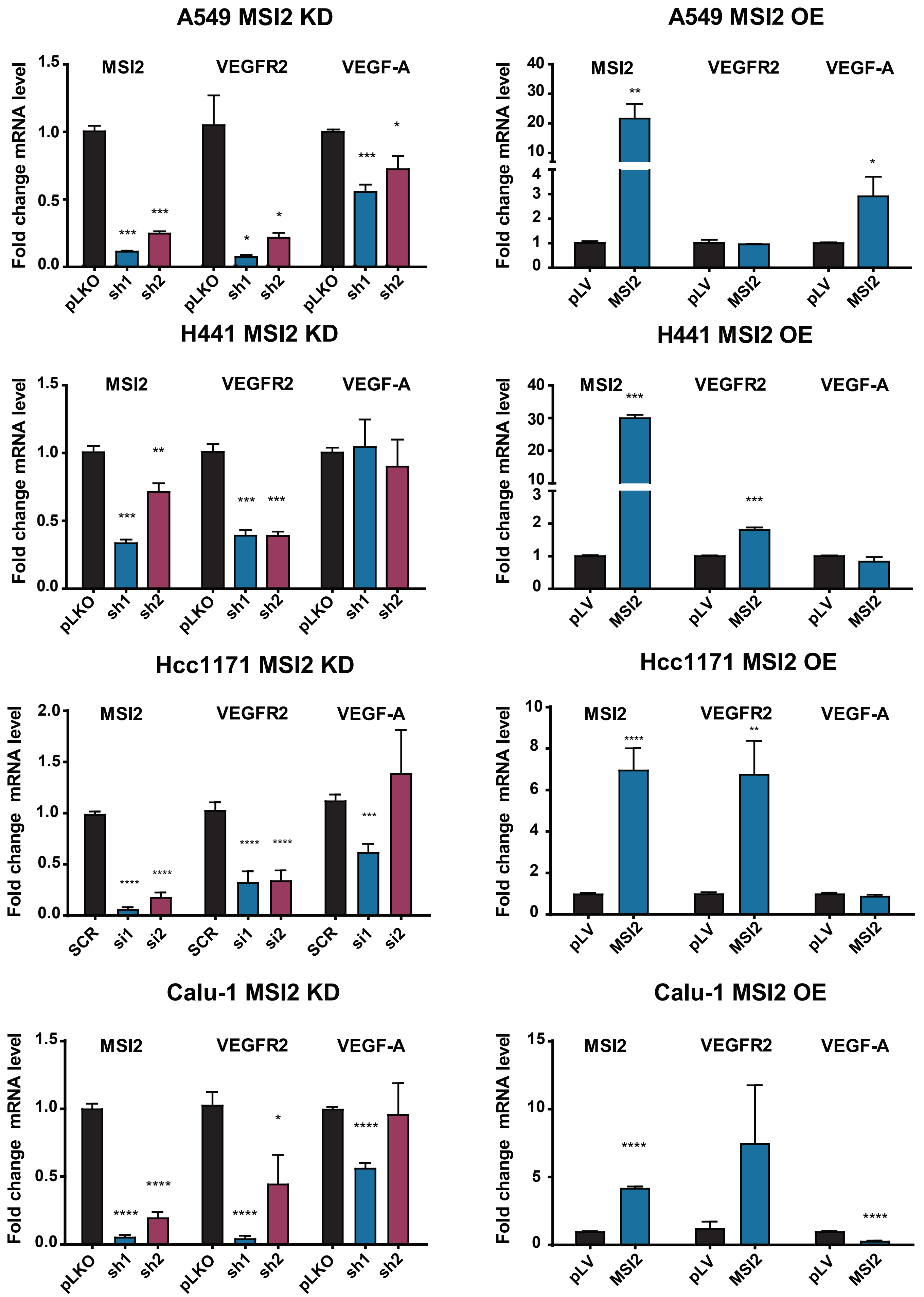


Supplementary Figure 3. Consequences of MSI2 depletion (left) and overexpression (right) on mRNA expression levels of VEGFR2 and VEGF-A. Quantitative RT-PCR of mRNA of human NSCLC cell lines, following MSI2 depletion by shRNA (sh1, sh2) and siRNA (si1, si2) and overexpression (MSI2) of MSI2. Negative controls include pLKO, SCR, pLV. Error bars represented by SEM. Statistical analysis was performed using an unpaired two-tailed t-test. *p < 0.05, **p < 0.01, ***p < 0.001.
