## Supplementary material for "Regulation of VEGFR2 and AKT signaling by Musashi-2 in lung cancer": Supp Table S2.docx

| Name | Vector | Type of insert | Type of expression | Cell line origin |
| --- | --- | --- | --- | --- |
| A549 pLKO | Tet-pLKO-puro | Empty | No | Human |
| A549 sh1 | Tet-pLKO-sh1-puro | MSI2-shRNA1 | Inducible | Human |
| A549 sh2 | Tet-pLKO-sh2-puro | MSI2-shRNA2 | Inducible | Human |
| A549 pLV | pLV-CMV-puro | Empty | No | Human |
| A549 MSI2 | pLV-CMV-MSI2-puro | MSI2 ORF | Constitutive | Human |
| A549 pHAGE | Tet-pLKO-puro  pHage-hygro | Empty  Empty | No  No | Human |
| A549 pHAGE MSI2 sh1 | Tet-pLKO-sh1-puro  pHage-hygro | MSI2-shRNA1  Empty | Inducible  No | Human |
| A549 VEGFR2 OE | Tet-pLKO-puro  pHage-KDR-hygro | Empty  KDR ORF | No  Constitutive | Human |
| A549 VEGFR2 OE MSI2 sh1 | Tet-pLKO-sh1-puro  pHage-KDR-hygro | MSI2-shRNA1  KDR ORF | Inducible  Constitutive | Human |
| H441 pLKO | Tet-pLKO-puro | Empty | No | Human |
| H441 sh1 | Tet-pLKO-sh1-puro | MSI2-shRNA1 | Inducible | Human |
| H441 sh2 | Tet-pLKO-sh2-puro | MSI2-shRNA2 | Inducible | Human |
| H441 pLV | pLV-CMV-puro | Empty | No | Human |
| H441 MSI2 | pLV-CMV-MSI2-puro | MSI2 ORF | Constitutive | Human |
| H441 pHAGE | Tet-pLKO-puro  pHage-hygro | Empty  Empty | No  No | Human |
| H441 pHAGE MSI2 sh1 | Tet-pLKO-sh1-puro  pHage-hygro | MSI2-shRNA1  Empty | Inducible  No | Human |
| H441 VEGFR2 OE | Tet-pLKO-puro  pHage-KDR-hygro | Empty  KDR ORF | No  Constitutive | Human |
| H441 VEGFR2 OE MSI2 sh1 | Tet-pLKO-sh1-puro  pHage-KDR-hygro | MSI2-shRNA1  KDR ORF | Inducible  Constitutive | Human |
| Calu-1 pLKO | Tet-pLKO-puro | Empty | No | Human |
| Calu-1 sh1 | Tet-pLKO-sh1-puro | MSI2-shRNA1 | Inducible | Human |
| Calu-1 sh2 | Tet-pLKO-sh2-puro | MSI2-shRNA2 | Inducible | Human |
| Calu-1 pLV | pLV-CMV-puro | Empty | No | Human |
| Calu-1 MSI2 | pLV-CMV-MSI2-puro | MSI2 ORF | Constitutive | Human |
| Hcc1171 pLV | pLV-CMV-puro | Empty | No | Human |
| Hcc1171 MSI2 | pLV-CMV-MSI2-puro | MSI2 ORF | Constitutive | Human |
| Hcc1171 pHAGE | pHage-hygro | Empty | No | Human |
| Hcc1171 VEGFR2 OE | pHage-KDR-hygro | KDR ORF | Constitutive | Human |
| 344SQ pLKO | pLKO.1-puro | Empty | No | Mouse |
| 344SQ sh1 | pLKO-m1-puro | MSI2-shRNA1 | Constitutive | Mouse |
| 344SQ sh2 | pLKO-m2-puro | MSI2-shRNA2 | Constitutive | Mouse |

**Supplementary table S2.** List of cell line derivatives used in the study. For human cell lines, the lentiviral vector Tet-pLKO-puro (Addgene, Plasmid #21915) was used for the inducible expression of shRNAs. For murine cell lines, the lentiviral vectors pLKO.1-puro (Addgene, Plasmid #8453) was used for the stable expression of shRNAs. For human cell lines, the pLV-CMV-puro vector (a kind gift from Dr. A. Ivanov, West Virginia University) was used for the stable expression of the MSI2 cDNA, the pHage-KDR-hygro vector was used for the stable expression of the VEGFR2 cDNA.
