## Supplementary material for "Regulation of VEGFR2 and AKT signaling by Musashi-2 in lung cancer": Supp Table S3.docx

| **siRNA Catalog Number** | **Gene Symbol** | **Sequence** |
| --- | --- | --- |
| SI04236652 and SI04285834 | Human MSI2 "-si1"  (Mixture 2) | ATGAGAGATCCCACTACGAAA and  CUGGAUUGGUCAUCAGAUU |
| SI04312665 and SI04375847 | Human MSI2 "-si2"  (Mixture 1) | TCCCAACTTCGTGGCGACCTA and CCAGATAGCCTTAGAGACTAT |

**Supplementary table S3. siRNAs targeting human and murine MSI2**. siRNA reference numbers are from Qiagen (Frederick, MD).
