## Supplementary material for "Regulation of VEGFR2 and AKT signaling by Musashi-2 in lung cancer": Supp Table S4.docx

| Gene symbol (H)-human; (M)-mouse | SYBR Green | Taqman Life Technologies |
| --- | --- | --- |
| *PTP4A1* (H) | Fw: 5`-ATCCAACCAATGCGACCTTA |  |
|  | Rev: 5`-AAGGCCAATCAAGAACATGG |  |
| *GAPDH* (H) | Fw: 5`-TGCACCACCAACTGCTTAGC |  |
|  | Rev: 5`-GGCATGGACTGTGGTCATGAG |  |
| *TGFBR1* (H) | Fw: 5`- ATCTTGTACCTTCTGACCCATC  Rev: 5`- TGGCATACCAACATTCTCTCAT |  |
| *VEGFR2* (H) | Fw: 5`-CAGAAATGTACCAGACCATGCT  Rev: 5`- GGAAGAACAATGTAGTCTTTGCC |  |
| *VEGF-A* (H) | Fw: 5`- CCATGAACTTTCTGCTGTCTTG |  |
|  | Rev: 5`- GCGCTGATAGACATCCATGA |  |
| *ERBB3* (H) | Fw: 5`-TGCTATACAGTGAGGCCAAGACTC |  |
|  | Rev: 5`-CAACTCCCAAACTGTCACACCATA |  |
| *SMAD3* (H) | Fw: 5`-CCAGCACATAATAACTTGGACCT |  |
|  | Rev: 5`-GATGTGTCTCCGTGTCAGCTC |  |
| *MSI2* (H) | Fw: 5`-GGTCATGAGAGATCCCACTACG  Rev: 5`-TCTACACTTGCTGGGTCTGC |  |
| *ACTB* (H) | Fw: 5`- TTGTTACAGGAAGTCCCTTGCC  Rev: 5`- ATGCTATCACCTCCCCTGTGTG |  |
| *PTEN* (H) | Fw: 5`- GCTCTATACTGCAAATGCTATCG  Rev: 5`- CCACAAACAGAACAAGATGCT |  |
| *18S rRNA* (H) |  | Fw: 5` GCTCTTTCTCGATTCCGT  Rev: 5`- CCAGAGTCTCGTTCGTTATC  Probe: 6fam- TTCTTAGTTGGTGGAGCGATTTGT- Iowa blackFQ |
| *Msi2* (M) | Fw: 5`-AGCAGTATTTCGAGCAGTTTGGCA  Rev: 5`-TGTCGAACATCAGCATCGCATCC |  |
| *Vegfr2* (M) | Fw: 5`-TTCAGAGTTGGTGGAGCATT  Rev: 5`-CTCTTCCATGCTCAGTGTCTC |  |
| *Vegfa* (M) | Fw: 5`-AGAAAGACAGAACAAAGCCAGA  Rev: 5`-TGGTGACATGGTTAATCGGT |  |
| *Polr2a*(M) | Fw: 5`- GGTCCTTCGAATCCGCATC  Rev: 5`- CAGGGTCATATCTGTCAGCATG |  |

**Supplementary Table S4**. **Primers used for RT-PCR to quantify gene expression.** List of primers used for SYBR Green assays, and TaqMan gene expression assays, used for the qRT-PCR analysis of gene expression and RNA immunoprecipitation.
