## Supplementary material for "Regulation of VEGFR2 and AKT signaling by Musashi-2 in lung cancer": Supplementary Figure 1.pdf

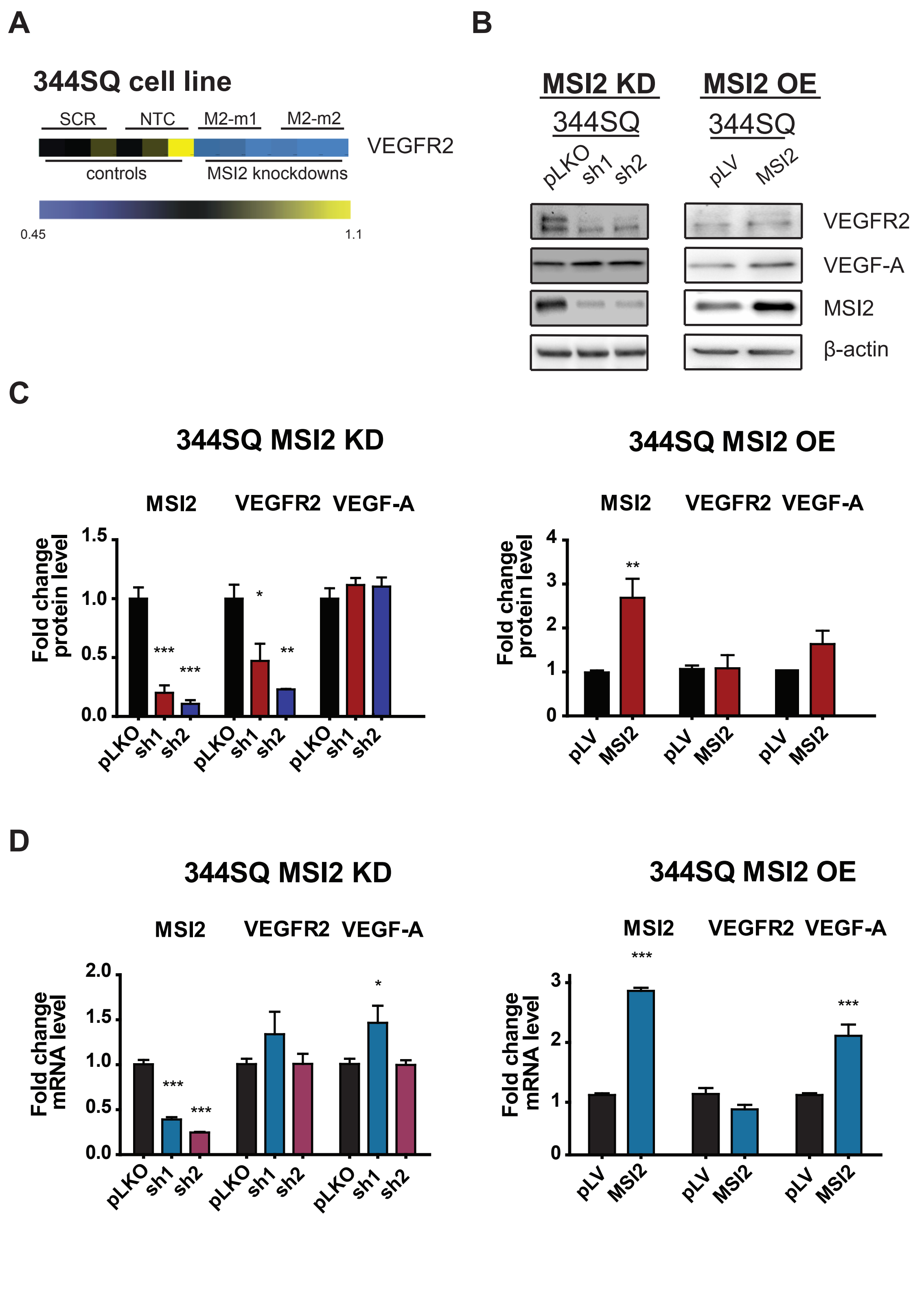


Supplementary Figure 1.
MSI2 regulation of VEGFR2 protein level in a mouse cell line. (A) Heatmap of RPPA results for VEGFR2 protein expression in 344SQ cell line transfected with empty vector or shRNAs or MSI2 depletion with two independent shRNAs (M2-m1 and M2-m2). (B) Western blots of 344SQ cell line lysates, following depletion (MSI2 KD sh1, sh2) and overexpression (MSI2 OE, MSI2) of MSI2. pLKO and pLV are negative controls. MSI2 depletion was induced by the addition of 1 µg/ml of Doxycycline for 48 h. (C) Quantification of Western blot data (Supplementary Fig. 1A) from at least three independent experiments by Image J software, with values normalized to negative control and β-actin. (D) RT-qPCR analysis of Vegfr2 and Vegf-a mRNA from 344SQ cell line following depletion (MSI2 KD, sh1, sh2) and overexpression (MSI2 OE, MSI2) of MSI2. pLKO and pLV are negative controls. MSI2 depletion was induced by the addition of 1 µg/ml of Doxycycline for 48 h. Data normalized to negative control and Polr2a. Error bars represented by SEM. Statistical analysis was performed using an unpaired two-tailed t-test. $^*p < 0.05$, $^{**}p < 0.01$, $^{***}p < 0.001$ for all graphs.
