## Supplementary material for "Regulation of VEGFR2 and AKT signaling by Musashi-2 in lung cancer": Supplementary Figure 4.pdf

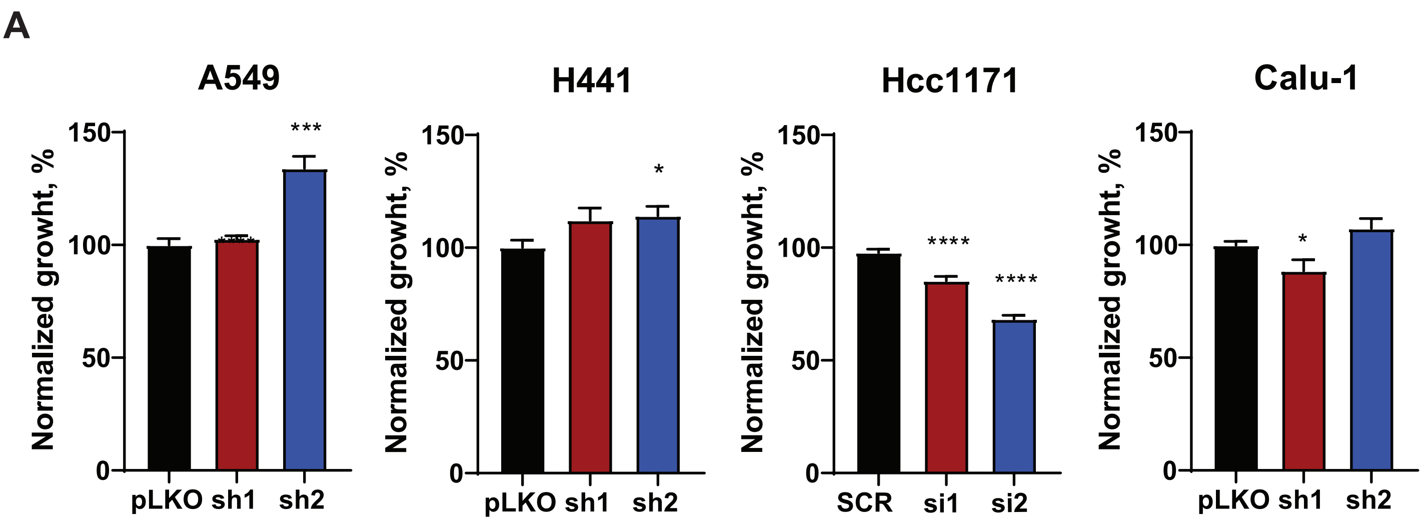


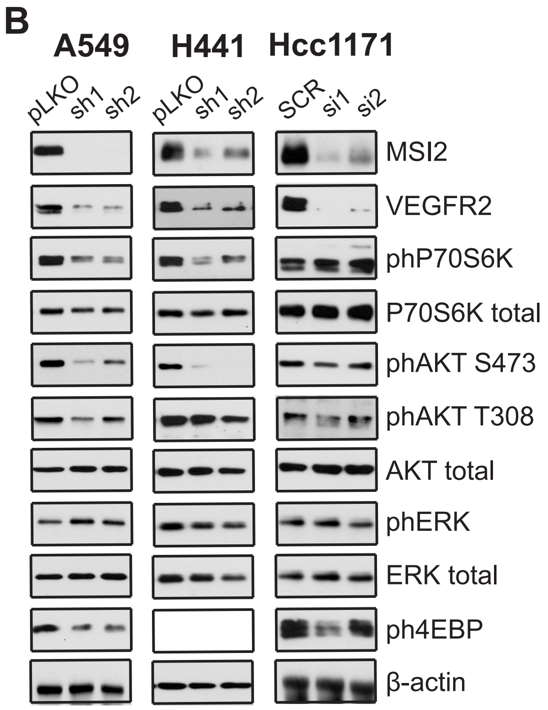


Supplementary Figure 4. (A) Cell viability quantified by Cell Titer Blue (CTB) assay of indicated cell lines following depletion by shRNA (sh1, sh2), siRNA (si1, si2) and overexpression (MSI2) of MSI2. MSI2 depletion was induced by the addition of 1 µg/ml of Doxycycline for 48h. Negative controls include pLKO or SCR. Error bars represented by SEM. Statistical analysis was performed using an unpaired two-tailed t-test. $*p < 0.05$, $**p < 0.01$, $***p < 0.001$, $****p < 0.0001$. (B) Western blots of indicated cell lines, following depletion by shRNA (sh1, sh2) and siRNA (si1, si2) of MSI2. Negative controls include pLKO and SCR. Error bars represented by SEM. Statistical analysis was performed using an ANOVA. $*p < 0.05$, $**p < 0.01$, $***p < 0.001$, $****p < 0.0001$.
